# Protein engineering shows antifreeze activity scales with ice-binding site area

**DOI:** 10.1101/2022.09.07.506985

**Authors:** Connor L. Scholl, Peter L. Davies

## Abstract

The ice-binding site (IBS) of the 9.6-kDa springtail (Collembola) antifreeze protein from *Granisotoma rainieri* was identified by mutagenesis. We then studied the protein’s activity as a function of IBS area. Its polyproline type II helical bundle fold facilitates changes to both IBS length and width. A one third increase in IBS width, through the addition of a single helix doubled antifreeze activity. A one third decrease in area reduced activity to 10%. A construct engineered with an additional tripeptide turn in each helix displayed a 5-fold decrease in activity. Molecular dynamics suggested that the lengthened IBS is more twisted than the wild type, emphasizing the importance of a flat surface for antifreeze activity.

## Introduction

Antifreeze proteins (AFPs) bind to, and prevent further growth of, ice crystals to help their hosts avoid freezing (1). Their activity can be evaluated by thermal hysteresis (TH), which is the difference between the elevated melting temperature and depressed freezing temperature (2). Binding of the AFP to the ice surface prevents further ice growth by making it energetically unfavourable for new water molecules to join the ice lattice (3). AFPs are proposed to bind to ice through waters that form organized structures around methyl groups on the ice-binding surface (IBS) and mimic waters in the quasi-liquid layer next to the ice lattice (4). These ice-like waters allow the protein to adsorb on, and become incorporated into, the ice surface (5). Surface water organization on the IBS has been observed in a few instances during crystallography (6-8), where crystallographic waters in these AFP structures resemble ice waters found in one or more planes of an ice crystal. However, most AFPs crystalize with their IBS facing one another due to the hydrophobicity of this surface, and this close contact tends to restrict the number of waters present. Yet, molecular dynamics simulations studying the binding mechanism of the beetle *Tenebrio molitor* AFP (*Tm*AFP) suggest that pre-ordered waters are not required for ice adsorption (9). Rather, slow diffusion of an AFP to the ice surface in the proper orientation may lead to rapid formation of clathrate waters. Thus, it is not clear if surface ice-like waters are the cause or effect of AFP binding to ice. As complete knowledge of the mechanism is lacking, additional functional studies are required to further understand this process.

When ice-binding proteins adsorb to the ice surface, they specifically bind to one or more ice planes, which can be defined by Miller indices (10). In the absence of AFP, ice will grow as a flat disk with only its basal plane well-defined. Upon binding, the AFP will structure the ice crystal into various hexagonal shapes depending on which plane it interacts with. AFPs that do not bind to the basal plane are typically moderately active, while AFPs that bind to more than one, one of which is the basal plane, tend to be hyperactive. The ability of hyperactive AFPs to bind to more than one ice plane improves its ability to further prevent ice growth. Moderately active AFPs have 0.5 - 1 °C TH activity at micromolar concentrations, while hyperactive AFPs have higher activity at lower concentrations (11).

In natural AFP isoforms, a correlation has been observed between the surface area of the IBS and the AFP’s antifreeze potency. It is common for AFP-producing organisms to produce multiple isoforms of AFPs that differ in size. For the alanine-rich type I AFP from winter flounder, the four-repeat isoform at 4 mg/mL had nearly double the antifreeze activity compared to that of the three-repeat isoform (12). An engineered *Tm*AFP showed a positive correlation between antifreeze potency and the number of β-helical loops each of which had a water organizing TxT motif (13). In these examples, the one-dimensional expansion of the repetitive region of alpha-helical AFPs or beta-solenoids led to non-linear increases in AFP potency.

The stand-alone polyproline type II (PPII) helix bundle fold used by springtail AFPs offers a chance to manipulate both the width and length of the IBS. Previously, a single layer of crystallographic waters on the proposed IBS of the 9.6-kDa springtail AFP from *Granisotoma rainieri* (*Gr*AFP) were shown to be organized like water molecules on both basal and primary prism planes of ice (14). The structure of *Gr*AFP is composed of nine antiparallel PPII helices that arrange themselves into two layers. The face of one layer is flat, contains small, hydrophobic residues, and is proposed to be the IBS. The opposing face is uneven with large residues some of which are charged. This structure is an ideal model to study how surface area affects TH activity as the IBS can be expanded two-dimensionally, through either duplication of the helices or elongation of the PPII helices.

Here, we looked to increase the area of IBS of *Gr*AFP to determine its affect on AFP activity. But first it was necessary to confirm the location of the proposed IBS by site-directed mutagenesis. A single lysine mutation on the proposed IBS decreased the antifreeze activity 10-fold, while two lysine mutations nearly eliminated antifreeze activity. A single lysine mutation on the opposing side presented similar activity as the wild type. Engineered constructs to alter the area of the IBS were designed by deleting or duplicating two central helices, resulting in one fewer (*Gr*AFP-7) or one greater helix (*Gr*AFP-11) on the IBS. With an additional ice-binding helix, the TH activity doubled, while removing an ice-binding helix lowered the activity to 10% relative to the wild type. In addition, an engineered construct with an additional turn in each helix was designed (*Gr*AFP-L). This elongated construct had a 5-fold decrease in activity compared to the wild type. Based on molecular dynamics, it is suggested that lengthening of the helix allows for more rotation along the axis of the helices. This torsion likely spoils the flatness of the IBS and its docking to ice.

## Materials and Methods

### Modelling and molecular dynamics

Engineered constructs were designed and visualized in PyMOL (Schrödinger, Inc., New York, NY, USA) using chain A of *Gr*AFP (7jjv) as a template. To confirm the stability of the constructs molecular dynamics using GROMACS (version 2016.3; University of Groningen, Groningen, Netherlands) was performed as previously described (14). Trajectories were visualized in VMD (version 1.9.3; University of Illinois, Champaign, IL, USA). The RMSD plots of the α-carbons were produced using R (R Core Team, 2021).

### Gene synthesis of engineered constructs

Codon-optimized, synthetic genes for *Gr*AFP-7, -11, and -L were ordered from GeneArt (ThermoFisher, Waltham, MA, USA). With the signal peptide of *Gr*AFP (QQY00623.1) removed, a methionine at the N terminus and leucine-glutamate at the C terminus were added, and residues were deleted or duplicated as follows: i) ***Gr*AFP-7**, residues 30 – 59 were deleted, ii) ***Gr*AFP-11**, residues 48 – 77 were duplicated, iii) ***Gr*AFP-L**, residues 8 – 10, 18 – 20, 31 – 33, 47 – 49, 63 – 65, 77 – 79, 92 – 94, 104 – 106 and 116 – 118 were duplicated, and S23A, V58A, and A93G were mutated. The genes were excised from the pMX vector, digested with *Nde*I and *Xho*I, and ligated into pET24-a(+) at those restriction sites, thus introducing a C-terminal His-tag. The ligated plasmids were then transformed into *E. coli* TOP10 competent cells (Invitrogen, Carlsbad, CA, USA). The plasmids were miniprepped and the DNA was sequenced. Once the sequence was confirmed, the plasmids were transformed into *E. coli* BL21 (DE3) expression cells (Invitrogen, Carlsbad, CA, USA).

### Site-directed mutagenesis of mutant constructs

From the previously synthesized *Gr*AFP gene (14), three primers were designed to mutate residues to identify the IBS. Each mutagenesis reaction was performed using QuikChange II Site-directed Mutagenesis kit (Agilent, Santa Clara, CA, USA) and the mixtures were made according to manufactures guidelines. The primers that were used are listed in **Table 1**. The reactions were heated to 95 °C for 2 min, then cycled through the following temperatures 16 times: 95 °C for 1 min, 55 °C for 1 min, 68 °C for 6 min. Once removed from heating, reaction tubes were cooled on ice to room temperature, then 10 U of *Dpn*I was added to each reaction. The reactions were mixed, then incubated at 37 °C for 30 min. The mutated plasmids were transformed into *E. coli* TOP10 competent cells (Invitrogen, Carlsbad, CA, USA). The plasmids were miniprepped and the DNA was sequenced. Once the sequences were confirmed, the plasmids were transformed into *E. coli* BL21 (DE3) expression cells (Invitrogen, Carlsbad, CA, USA).

**Table 1.**
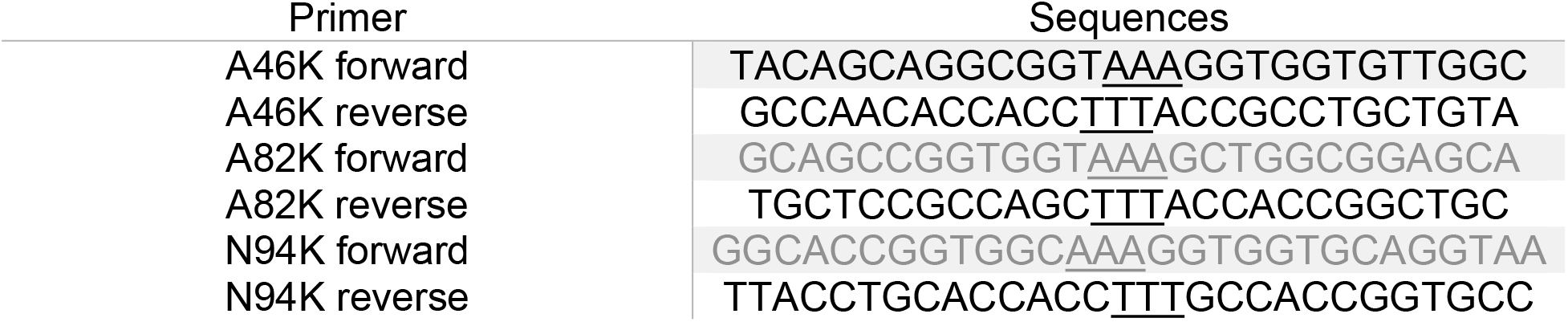
*Gr*AFP mutant construct primers.

### Recombinant expression and purification

Cells plated on kanamycin-resistant agar plates were picked into flasks containing 100 mL lysogeny broth (LB) media with 100 µg/mL kanamycin. The inoculated cultures were incubated shaking at 37 °C overnight. The overnight cultures were divided eventually among four flasks containing 1 L LB with 100 µg/mL kanamycin. The flasks were then incubated shaking at 37°C until the OD_600_ was 0.6 – 1.0. Once cooled to room temperature, 1 mM isopropyl ß-D-1-thiogalactopyranoside was added to each culture and the cells were induced during overnight shaking at 20 °C. The cells were collected by centrifugation and were resuspended in 50 mL of resuspension buffer (20 mM Tris/HCl (pH 7.8), 500 mM NaCl, 5 mM imidazole, and one cOmplete™ ultra protease inhibitor cocktail tablet (Roche Applied Sciences, Mannheim, Germany)). The cells were lysed with 16 × 10 s rounds of sonication with cooling below 6 °C between rounds to prevent thermal denaturation. The lysate supernatants were mixed with 5 mL of HisPure Ni-NTA resin for 30 min at 4 °C. The mixtures were then added to a gravity column and washed with 50 mL of wash buffer (20 mM Tris/HCl (pH 7.8), 500 mM NaCl, and 10 mM imidazole). The His-tagged protein was eluted with 5 × 5 mL of elution buffer (20 mM Tris/HCl (pH 7.8), 200 mM NaCl, 200 mM imidazole). The presence of target proteins in the elutions were confirmed with sodium dodecyl sulfate-polyacrylamide gel electrophoresis, the positive fractions were pooled, and the volume was brought to 50 mL with buffer (20 mM Tris/HCl (pH 7.8) and 200 mM NaCl). The pooled solutions were added to round bottom flasks seeded with 10 – 15 mL of ice and the round-bottom flasks were rotated in baths with cooled ethylene glycol to -1.3 °C for 90 min (15). The temperatures were then lowered by 0.2 °C every 30 min until 50% of the samples’ volumes were incorporated into the ice-shells. After the liquid fractions were decanted, the ice fractions were melted with 5 mL of 200 mM NH_4_HCO_3_ and the volumes were increased to 50 mL with ddH_2_O. The samples were concentrated using an Amicon Ultra 3K Centrifugal Filter (MilliporeSigma, Burlington, MA, USA) spun at 4 °C in a Sorvall ST16R centrifuge at 3000 x *g*. The samples were buffer exchanged into 20 mM NH_4_HCO_3_ using the same filter. The concentrations of the final purified samples were calculated using acid hydrolysis amino acid analysis (SPARC, The Hospital for Sick Children, Toronto, ON, Canada). The samples were then aliquoted, flash-frozen with liquid nitrogen, and stored at -80 °C.

### Thermal hysteresis

The thermal hysteresis measurements were performed as previously described (16). The samples were injected into an oil droplet-containing grid place on a Peltier unit. The temperatures were controlled using a model 3040 temperature controller (Newport, Irvine, CA, USA) and cooled with a Clifton nanoliter osmometer. The oil-submerged protein samples were flash-frozen, then slowly melted until a single ice crystal remained. The temperature was then decreased at a rate of -0.005 °C per 4 s until ice growth commenced. Videos of the ice crystals were recorded using a DMK 33UX249 USB 3.0 monochrome industrial camera (The Imaging Source, Charlotte, NC, USA).

## Results

### Site-directed mutagenesis identifies the ice-binding surface of *Gr*AFP

Mutations were made to the two flat surfaces of *Gr*AFP (**Fig. 1A**) to determine the location of the IBS. The mutant construct A46K inserted a lysine residue on helix 4 that added a bulky, charged residue onto the originally flat, hydrophobic face that was predicted to be the IBS (**Fig. 1B**). An additional lysine residue was added on the flat surface on helix 6, which produced the double mutant *Gr*AFP A46K A82K (**Fig. 1C**). To establish that adding a lysine residue elsewhere does not cause a change in TH activity, a mutant N94K with a lysine residue in the opposing surface was produced (**Fig. 1D**). The ice-shaping and ice growth patterns were altered with the additions of lysine residues on the IBS of *Gr*AFP (**Fig. 1E**). For the wild type, the ice crystal began as an oval-shaped crystal but as the temperature decreased it took on a hexagonal bipyramidal shape. At the freezing point, the ice crystal had a rapid “burst” with ice spicules emanating along the crystal’s *a*-axes. This type of ice-shaping and burst pattern was also seen in the N94K mutant (not shown). With the addition of a single lysine residue on the IBS, *Gr*AFP A46K had a reduced ability to protect the growth of the basal plane. The ice crystal presented as a hexagonal column with bipyramid tips but as the temperature was decreased the ice crystal began growing parallel to the *c*-axis more rapidly than the other planes. The double mutant, A46K A82K, had a more truncated hexagonal bipyramidal shape and grew out of all planes at a similar rate.

**Figure 1.**
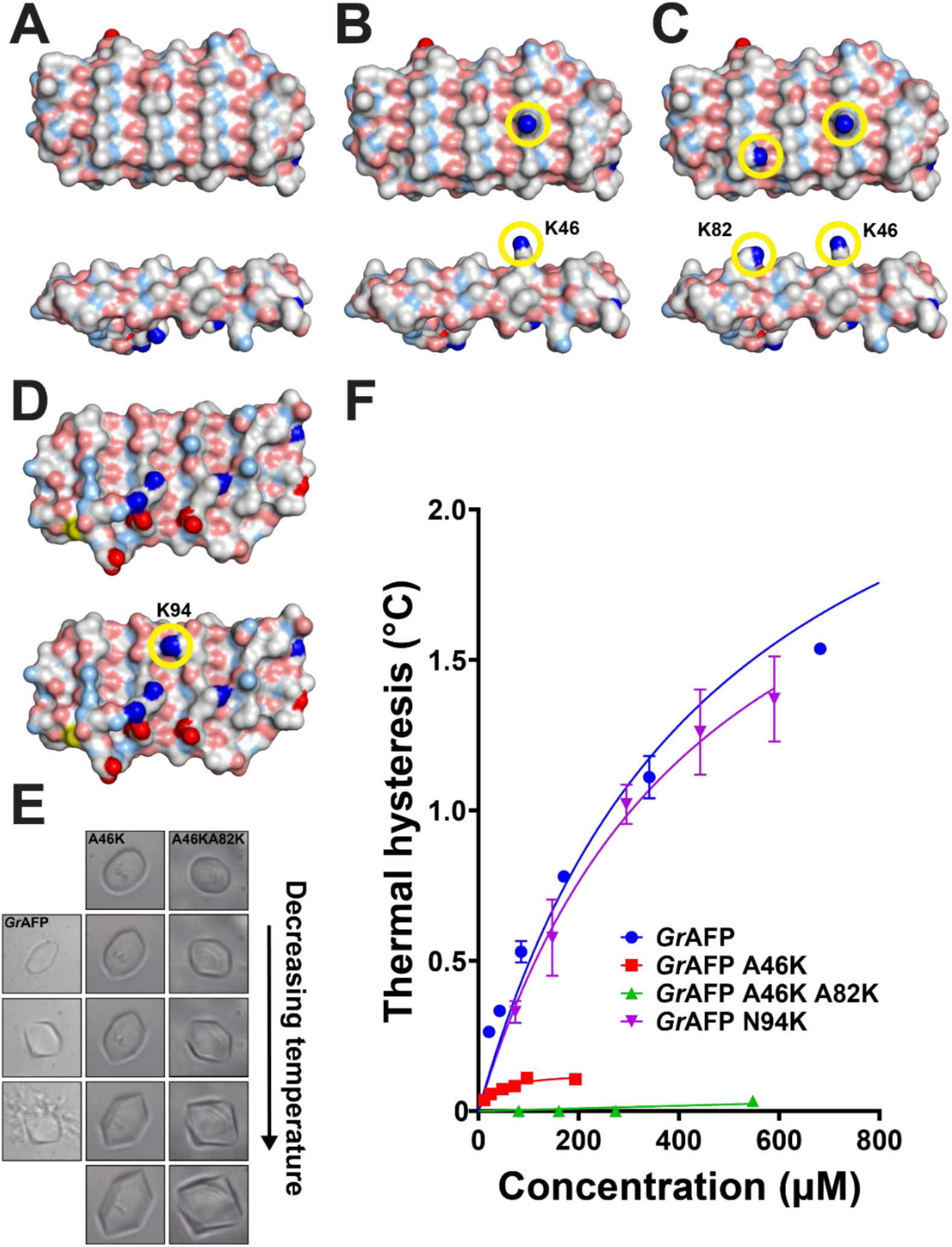
*Gr*AFP ice-binding site identification. **A-C)** Top-down (above) and side-on (below) views of putative ice-binding surface of **A)** *Gr*AFP, **B)** *Gr*AFP A46K and **C)** *Gr*AFP A46K A82K. Mutated residues are highlighted with a yellow circle. **D)** Top-down view of the non-ice-binding surface of *Gr*AFP (top) and *Gr*AFP N94K (bottom). **E)** The ice-shaping progression of a single crystal in solution with *Gr*AFP, *Gr*AFP A46K, or *Gr*AFP A46K A82K. Panels going top to bottom with decreasing temperature until ice growth occurs. **F)** Thermal hysteresis curves of purified *Gr*AFP (blue circles), *Gr*AFP A46K (red squares), *Gr*AFP A46K A82K (green triangles), and *Gr*AFP N94K (purple triangles).

The thermal hysteresis activities of the mutants were compared to the wild type (**Fig. 1F**). For the single lysine mutant, the TH curve had slight initial increase but soon plateaued reaching a TH of ∼0.2 °C. Compared to the wild type, *Gr*AFP A46K had greatly reduce antifreeze activity, having about 10% activity of that of the wild type. The A46K A82K mutant had a near-linear TH curve and showed little TH activity with just 0.03 °C at 555 µM. Interestingly, despite their lower TH activities, both single and double lysine mutants were isolated using ice-affinity purification.

The N94K mutant had similar activity to the wild type with the TH curve having an initial steep increase, as seen in the wild type. Due to the poorer solubility of *Gr*AFP N94K, the concentration was only a third of that of the wild type, so the maximum TH, or TH_max_, could only been estimated based on extrapolating the curve.

### Lengthening the polyproline type II helical AFP did not increase its activity

The consequences of elongating the PPII helices were explored. A single GXX peptide repeat was duplicated in each of the helices, increasing the number of ice-ordering repeats in *Gr*AFP (**Fig. 2A**) from three turns to four. To compare the IBS length, the distance between the first and last alanine residues on the PPII helix was used. Thus, the IBS of the lengthened *Gr*AFP (*Gr*AFP-L) has an increased surface area of ∼33% relative to *Gr*AFP (**Fig. 2B**). During molecular dynamics analysis of the model there was noticeable oscillation in the RMSD of *Gr*AFP-L. The mean RMSD of *Gr*AFP (**Fig. 2C**; 1.06 ± 0.23 Å) was 0.36 Å lower than that of *Gr*AFP-L (**Fig. 2D**; 1.42 ± 0.25 Å).

**Figure 2.**
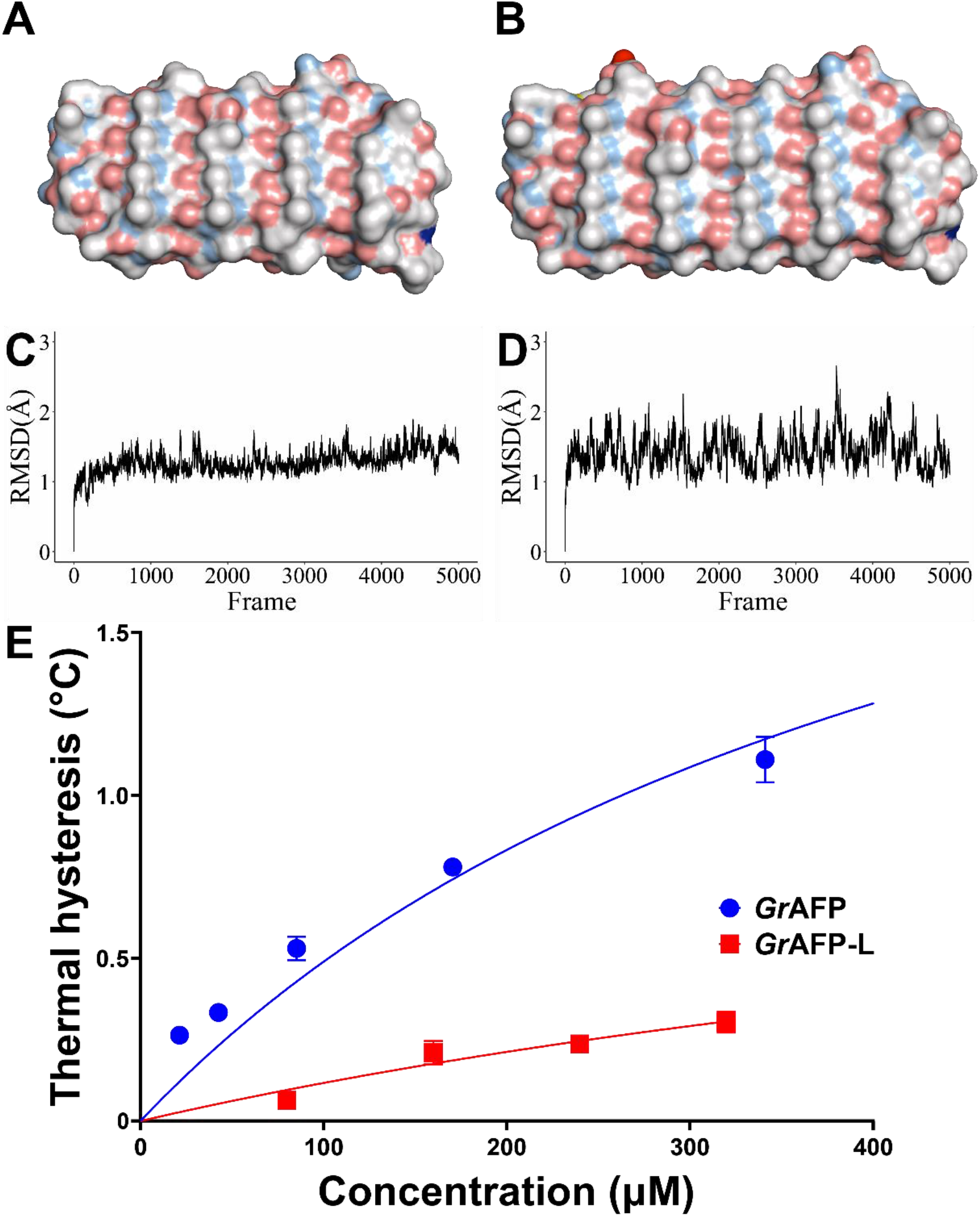
Characterization of *Gr*AFP helix elongation. The structure of **A)** *Gr*AFP was lengthened by one helical turn in each of the nine polyproline type II helices to design **B)** *Gr*AFP-L. The α-carbon RMSD of **C)** *Gr*AFP and **D)** *Gr*AFP-L monitored fluctuations are displayed during the 20-ns molecular dynamics analyses. **E)** The thermal hysteresis curves of purified *Gr*AFP (blue circles) and *Gr*AFP-L (red squares) as a function of protein concentration.

The TH curve of *Gr*AFP-L shows that it has less potent antifreeze activity compared to *Gr*AFP (**Fig. 2E**). The fitted trendline is slightly curved and does not appear to reach a maximum with the plotted data. As was the case with *Gr*AFP-11, the increased surface area of the hydrophobic IBS reduced the protein’s solubility. Despite having an increased IBS surface area, extrapolation of the fitted curve estimates the TH_max_ to be ∼0.5 °C, a 5-fold decrease with respect to *Gr*AFP (2.5 °C).

### Design process of *Gr*AFP engineered constructs

To produce the engineered *Gr*AFP constructs with helical deletions or duplications, models based on the structure of *Gr*AFP (**Fig. 3A**) were first designed to ensure the deletions would not alter the overall fold of the protein. Since helices one, two, and nine are outwards facing and contain GXX repeats, these helices were not optimal for deletion or duplication. Rather, the central helices consisting of GGX repeats were selected due to their inward-facing glycine residues, as these are essential for the internal hydrogen bonding network that stabilizes this fold. Given that the helices have less structural variation than the loop regions, residues were selected starting and ending from the center of the helices. This allowed the protein chains to be reattached following the removal or additional helices in the models with ease. The following deletions or duplications were made: i) GrAFP-7, residues 30 (helix three) to 59 (helix five) were removed (**Fig. 3B**); ii) *Gr*AFP-11 residues 48 (helix four) to 77 (helix six) were duplicated (**Fig. 3C**); and iii) *Gr*AFP-13 residues 48 (helix four) to 104 (helix eight) were duplicated (**Fig. 3D**). Energy minimization simulations using GROMACS were performed to assess the stability of the models. The constructs, similar to the wild type, remained stable throughout the simulation with stabilized RMSD values through 20 ns (**Fig. 3E**). These engineered constructs were then recombinantly expressed and purified (**Fig. 3F**).

**Figure 3.**
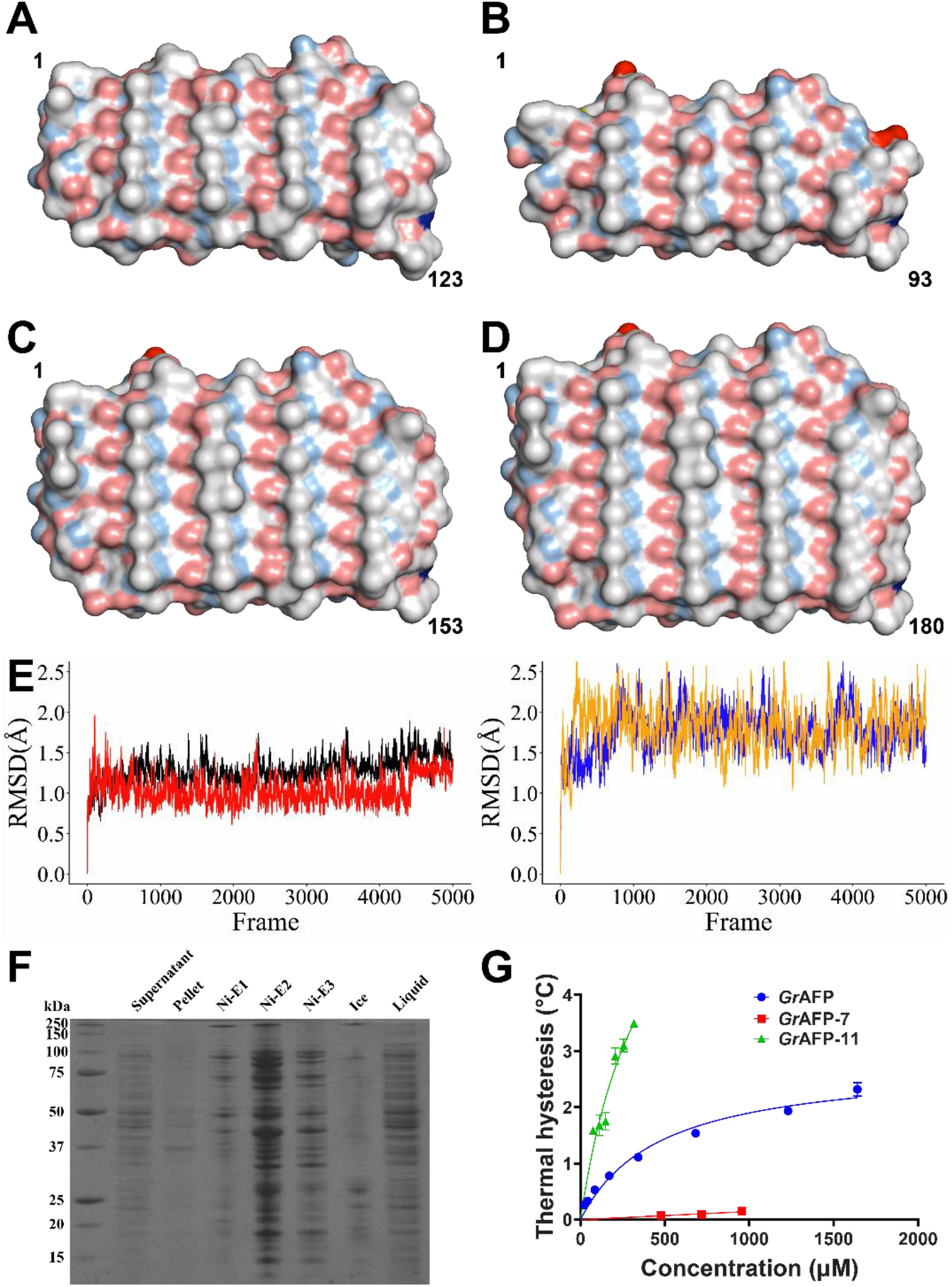
Characterization of polyproline type II helix deletion and duplication. The structure of **A)** *Gr*AFP was modified through deletion or duplication. **B)** *Gr*AFP-7 was modelled representing deletion of two helices, and **C)** *Gr*AFP-11 and **D)** *Gr*AFP-13 were modelled to show the duplication of two and four helices, respectively. The residue numbers are denoted at each terminus. **E)** The α-carbon RMSD of *Gr*AFP (black), *Gr*AFP-7 (red), *Gr*AFP-11 (blue), and *Gr*AFP-13 (orange) monitored across 20 ns of molecular dynamics simulations. **F)** Sample SDS-PAGE gel for the purification of *Gr*AFP-11. **G)** Thermal hysteresis of purified *Gr*AFP (blue circles), *Gr*AFP-7 (red squares), and *Gr*AFP-11 (green triangle) as a function of protein concentration.

### Helical deletion and duplication have consequences for the activity of *Gr*AFP

To determine the effect removing or adding helices had on the TH activity of *Gr*AFP, the activity vs. concentration of the purified proteins were compared (**Fig. 3G**). The removal of a single ice-binding helix in *Gr*AFP-7 had a negative effect on the maximum TH activity, decreasing it by over 10-fold (0.20 °C) compared to the wild type (2.5 °C). The activity-concentration relationship also appeared to be near-linear, and this contrasted to the hyperbolic curve seen in *Gr*AFP.

The addition of a single ice-binding helix in *Gr*AFP-11 had a non-linear effect on the TH activity, increasing it 3-fold (3.5 °C) at 315 µM compared to the wild type (∼1 °C). With this extra ice-binding helix, the overall solubility and expression with respect to *Gr*AFP decreased as the protein could not be concentrated past 315 µM. Even during concentration, shorter spin cycles were required to prevent local accumulation and precipitation to further increase the concentration. Due to this effect, the TH curve does not reach a plateau, but extrapolation of the curve estimated TH_max_ to be ∼5.5 °C. Following this trend, the addition of another ice-binding helix to form *Gr*AFP-13 produced trace amounts of soluble protein with much of the expressed protein found in inclusion bodies. Attempts to denature and refold the protein by dropwise addition to a renaturation buffer proved futile. Additional constructs with an engineered disulfide bridge on either the N- or C-terminal helix to aid in folding were also unsuccessful.

## Discussion

It is well established that the IBS of an AFP is flat and relatively hydrophobic, properties that have been supported through the mutation of putative water-ordering residues (17-19). In this study on *Gr*AFP the mutation of a single alanine to lysine residue decreased TH_max_ from ∼2.5 °C to ∼0.2 °C. The substitution of an alanine residue by a lysine residue on the IBS likely affected its function in two ways: 1) the replacement of the methyl group may have disrupted water clathrate formation in its vicinity; and 2) the amino group on the lysyl sidechain may have spoiled the hydrophobicity of the IBS that is needed for clathrate water formation. Similar trends have been seen in the β-helical AFPs from beetles *Tenebrio molitor* (*Tm*AFP) and *Rhagium mordax* (*Rm*AFP). In *Tm*AFP, replacement of the water-ordering threonine residues had a deleterious effect on the TH activity (20). Conversely, *Rm*AFP has a histidine and a lysine residue on the periphery of its IBS. When these were mutated to threonine residues, there was an increase of the TH activity (21). In contrast, the introduction of a lysine residue (N94K) on the non-ice binding face of *Gr*AFP had little effect on TH activity.

Analysis of ice-shaping can also be useful for qualitatively assessing ice-binding capabilities (22). Compared to *Gr*AFP, A46K and A46K + A82K cause changes in ice-shaping, whereas N94K does not. Also, the change in ice-shaping in going from zero to one to two lysine insertions suggests that the mutations weaken affinity for both the basal and primary prism planes of ice. A qualitative loss of ice shaping was also seen in the carrot ice-binding protein, where mutation of the glutamine and asparagine on the IBS led to less defined ice shaping (23).

This is the first AFP model that has the capability of being expanded two-dimensionally, as the IBS can be enlarged by both adding and elongating PPII helices. Indeed, the repetitive nature of the PPII helical bundle ice-ordering motifs allowed the protein to be engineered with ease. By adding a tripeptide GXX repeat to all helices, the ice-binding surface area increased by approximately 33%, as did the number of water-ordering motifs (12 to 16 motifs). This increase in area is identical to adding a single ice-binding helix on the IBS. However, the same increase in activity was not observed. With elongation, *Gr*AFP-L has a third the activity of *Gr*AFP at relative concentrations. Not only is the surface area and number of water-ordering residues important, but the spacing among these residues is also vital. When three or four β-coils were added to *Tm*AFP, a decrease in TH activity was observed from the +2-coil construct, albeit not lower than that of the wild type (24). The slight angle mismatches between threonine residues on adjacent β-strands caused a slight bend on the IBS. The addition of the third and fourth coils added no further contribution to binding. In the crystal structure of *Gr*AFP (7jjv), there is no curvature along the IBS. In contrast, during molecular dynamics, the surface begins to slightly twist along the helices. The elongation of the helices potentially allows for greater rotation, leading to a misalignment of the ice-like waters. Amongst the PPII helical bundle AFPs from Collembola, all known structures and models consist of three repeating units along the length of the helix(14,25-27). This is a potential reason why this protein family exists solely with three repeats. However, it should be noted that isoforms that arise through duplication events are more likely to have helical duplications rather than elongations. It would only take a single duplication for *Gr*AFP to have an additional ice-binding helix; however, in *Gr*AFP it would take nine duplication events for all helices to become elongated.

The strategy of duplicating existing helices proved effective, as the existing dihedral angles of the PPII helix were preserved. Helices were chosen to be duplicated in pairs due to their arrangements, with the IBS having four helices and the opposite face having five helices. If a single helix were added, it would be problematic in the following ways: 1) a single helical addition would alter the overall symmetry of the protein; and 2) the single helical addition would have to be on either the N or C terminus. This is troublesome as the N- and C-terminal helices contain GXX repeats rather than GGX. Additionally, the N- and C-terminal helices have greater variability compared to the interior helices, and this is accommodated on the N terminus with a disulphide bridge. Although these issues could have been overcome through more extensive engineering, it was simpler to duplicate interior helices in pairs.

The IBS should be looked at in two ways: 1) total surface area; and 2) number of ice-ordering motifs. Measuring the dimensions of the protein defines the surface area; however, one needs to consider if area scales with the number of ice-ordering motifs. When looking at the crystallographic waters in the structure of *Gr*AFP (7jjv), water molecules are found between each of the R-groups on the IBS. With four R-groups on each helix, this gives three water-ordering motifs per helix. The removal of a single ice-binding helix decreased the surface area by roughly 33% while the number of ice-ordering motifs decreased by 25% (12 to 9 motifs) and led to a 10-fold decrease in activity. With an additional ice-binding helix, there was approximately a 25% increase in surface area and a 33% increase in water-ordering motifs, and the potency increased by 2-fold. In this case, the TH activity changed non-linearly relative to the surface area and the number of ice-binding motifs. This trend has been seen with other AFP folds, although this is the first time it has been seen with the PPII helix bundle fold. The trend is evident with linear AFPs, such as in the type I alanine-rich AFPs from the winter flounder, where AFP9 has four repeating units while HPLC-6 has three repeating units (33% larger). Although the TH_max_ of AFP9 (∼1.0 °C) is only 25% higher than HPLC-6 (0.8 °C), it has a much steeper curve having nearly double the activity at 4 mg/mL (12). This surface area relationship has also been observed in β-solenoid AFPs from spruce budworm and *T. molitor*. The Spruce budworm isoform CfAFP-337 has two fewer coils than CfAFP-501 with a surface area difference of approximately 33% and 40% fewer ice-binding motifs (28). The seven-coil *Tm*AFP was engineered to have 6 - 11 coils and the activity increased with a growing number of coils (24). Removal of a single coil (15% less surface area) decreased the activity 5-fold, while the addition of a coil (14% greater surface area) increased the activity 2-fold.

We have shown that there is a positive correlation between the surface area of *Gr*AFP and its antifreeze activity. A small increase in area produces a disproportionately large increase in activity. Similarly, a small decrease in area produces a disproportionately large decrease in activity. However, there do seem be worsening solubility issues as the hydrophobic face of the PPII bundle increases in area in going from *Gr*AFP-9 to *Gr*AFP-11 to *Gr*AFP-13. It will be essential to overcome the solubility issue to explore the upper limits of the activity gain with increased IBS surface area.

Looking at natural occurring isoforms may suggest a solution. For example, in the springtail AFP from *Hypogastrura harveyi*, the 15.7-kDa AFP had twice the TH activity of the 6.5-kDa isoform (9). This protein has 11 PPII helices in the proposed model but is still relatively soluble. In a different IBP example, *Rm*AFP has histidine and lysine residues on the periphery that might help increase its solubility. Pushing the upper limit of AFP engineering will give a broader understanding of their mechanisms and allow for future biotechnological applications such as frost resistance and cryopreservation.

## Author Contributions

CLS and PLD initiated the research project; CLS performed experiments; CLS and PLD analyzed data; CLS drafted the manuscript; CLS and PLD edited the manuscript and approved its submission.

## Acknowledgements

We thank Dr. Robert Campbell for assistance with molecular dynamics and for his PyMOL scripts. This research was supported by CIHR Foundation Grant 148422 awarded to PLD. PLD holds the Canada Research Chair in Protein Engineering.

## Abbreviations

AFP: Antifreeze protein
TH: thermal hysteresis
IBS: ice-binding surface
*Tm*AFP: *Tenebrio molitor* antifreeze protein
PPII: polyproline type II
*Gr*AFP: *Granisotoma rainieri* antifreeze protein
LB: lysogeny broth
*Rm*AFP: *Rhagium mordax* antifreeze protein

